# The variant call format provides efficient and robust storage of GWAS summary statistics

**DOI:** 10.1101/2020.05.29.115824

**Authors:** Matthew Lyon, Shea J Andrews, Ben Elsworth, Tom R Gaunt, Gibran Hemani, Edoardo Marcora

**Author notes:** These authors contributed equally to this work (Matthew Lyon, Shea J Andrews). These authors contributed equally to this work (Gibran Hemani, Edoardo Marcora). **Correspondence** Matthew Lyon Population Health Sciences Bristol Medical School University of Bristol Oakfield House Oakfield Grove Bristol BS8 2BN.

## Abstract

Genome-wide association study (GWAS) summary statistics are a fundamental resource for a variety of research applications ^1–6^. Yet despite their widespread utility, no common storage format has been widely adopted, hindering tool development and data sharing, analysis and integration. Existing tabular formats ^7,8^ often ambiguously or incompletely store information about genetic variants and their associations, and also lack essential metadata increasing the possibility of errors in data interpretation and post-GWAS analyses. Additionally, data in these formats are typically not indexed, requiring the whole file to be read which is computationally inefficient. To address these issues, we propose an adaptation of the variant call format^9^ (GWAS-VCF) and have produced a suite of open-source tools for using this format in downstream analyses. Simulation studies determine GWAS-VCF is 9-46x faster than tabular alternatives when extracting variant(s) by genomic position. Our results demonstrate the GWAS-VCF provides a robust and performant solution for sharing, analysis and integration of GWAS data. We provide open access to over 10,000 complete GWAS summary datasets converted to this format (available from: https://gwas.mrcieu.ac.uk).

## Main

The GWAS is a powerful tool for identifying genetic loci associated with any trait, including diseases and clinical biomarkers, as well as non-clinical and molecular phenotypes such as height and gene expression ^3^ (eQTLs). Sharing of GWAS results as summary statistics (i.e. variant, effect size, standard error, p-value etc.) has enabled a range of important secondary research applications including: causal gene and functional variant prioritisation ^1^, causal cell/tissue type nomination ^2^, pathway analysis ^3^, causal inference (Mendelian randomization; MR) ^4^, risk prediction ^3^, genetic correlation ^5^ and heritability estimation ^6^. However, the utility of GWAS summary statistics is hampered by the absence of a universally adopted storage format and associated tools.

Historic lack of a common standard has resulted in GWAS analysis tools outputting summary statistics in different tabular formats (e.g. plink ^10^, GCTA ^11^, BOLT-LMM ^12^, GEMMA ^13^, Matrix eQTL ^14^ and meta-analysis tools e.g. METAL ^15^). As a consequence, various processing issues are typically encountered during secondary analysis. First, there is often inconsistency and ambiguity of which allele relates to the effect size estimate (the “effect” allele). Confusion over the effect allele can have disastrous consequences on the interpretation of GWAS findings and the validity of post-GWAS analyses. For example MR studies may provide causal estimates with incorrect effect directionality ^16^. Likewise, prediction models based on polygenic risk scores might predict disease wrongly or suffer reduced power if some of the effect directionalities are incorrect. Second, the schema (i.e. which columns/fields are included and how they are named) of these tabular formats varies greatly. Absent fields can limit analyses and although approaches exist to estimate the values of some of these missing columns (e.g. standard error from P value) imprecision is introduced reducing subsequent test power. Varying field names are easily addressed in principle, but the process can be cumbersome and error prone. Third, data are frequently distributed with no or insufficient metadata describing the study, trait(s), and variants (e.g., trait measurement units, variant id/annotation sources, etc.) which can lead to errors, impede integration of results from different studies and hamper reproducibility. Fourth, querying unindexed text files is slow and memory inefficient, making some potential applications computationally infeasible (e.g. systematic hypothesis-free analyses).

Some proposals for a standard tabular format have been made. The EBI-NHGRI GWAS catalog (www.ebi.ac.uk/gwas) developed a tab-separated values (TSV) text format with a minimal set of required (and optional) columns along with standardised headings ^7^. The SMR tool ^8^ introduced a binary format for rapid querying of quantitative trait loci. These approaches are adequate for storing variant level summary statistics but do not enforce allele consistency or support embedding of essential metadata. Learning from these examples and our experiences performing high-throughput analyses across two research centres, we developed a set of requirements for a suitable universal format (Table 1). These features place emphasis on consistency and robustness, capacity for metadata to provide a full audit trail, efficient querying and file storage, ensuring data integrity, interoperability with existing open-source tools and across multiple datasets to support data sharing and integration. We determined that adapting the variant call format (VCF) ^9^ was a convenient and constructive solution to address these issues. We provide evidence demonstrating how the VCF meets our requirements and showcase the capabilities of this medium (Table 1).

**Table 1.**
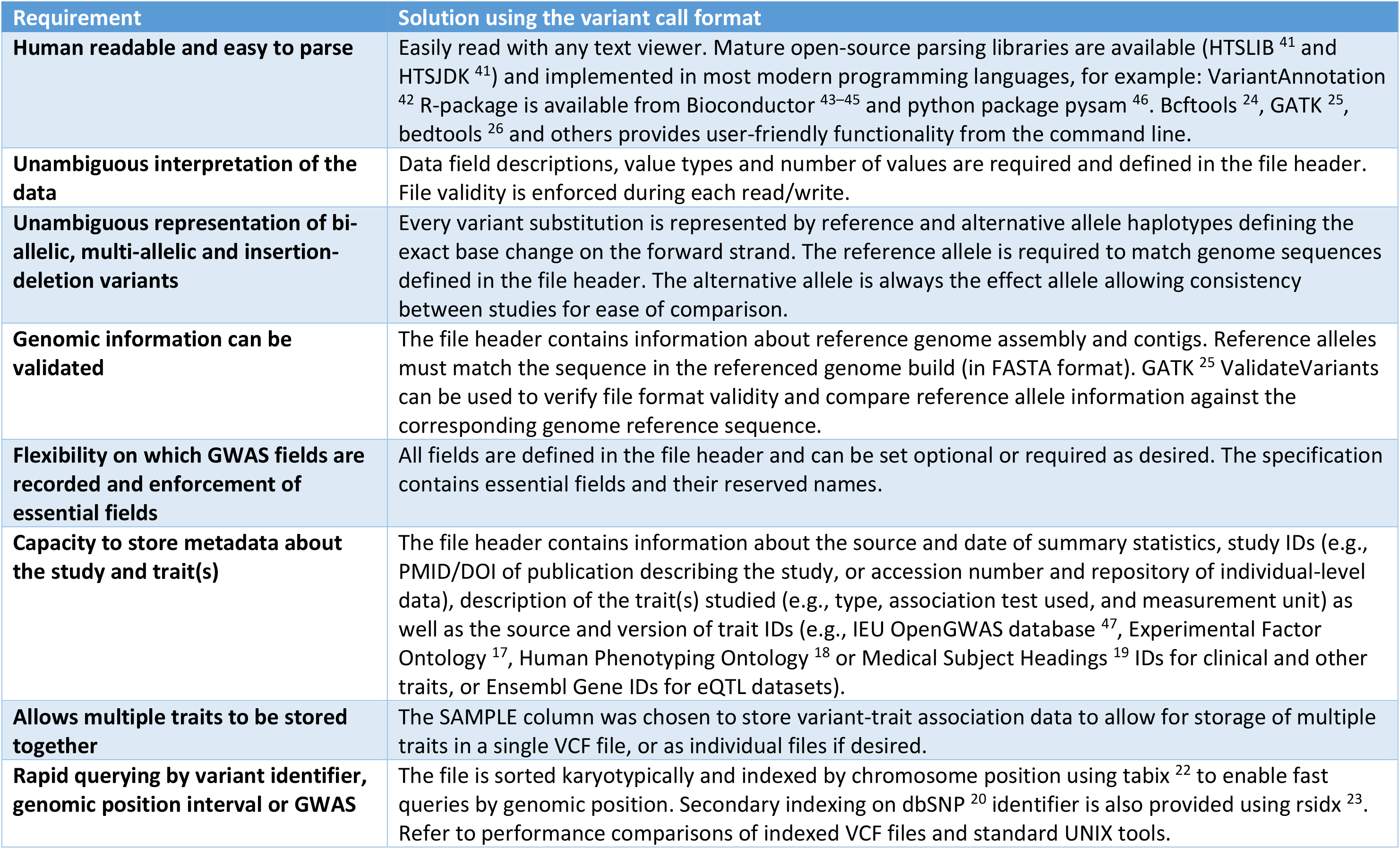
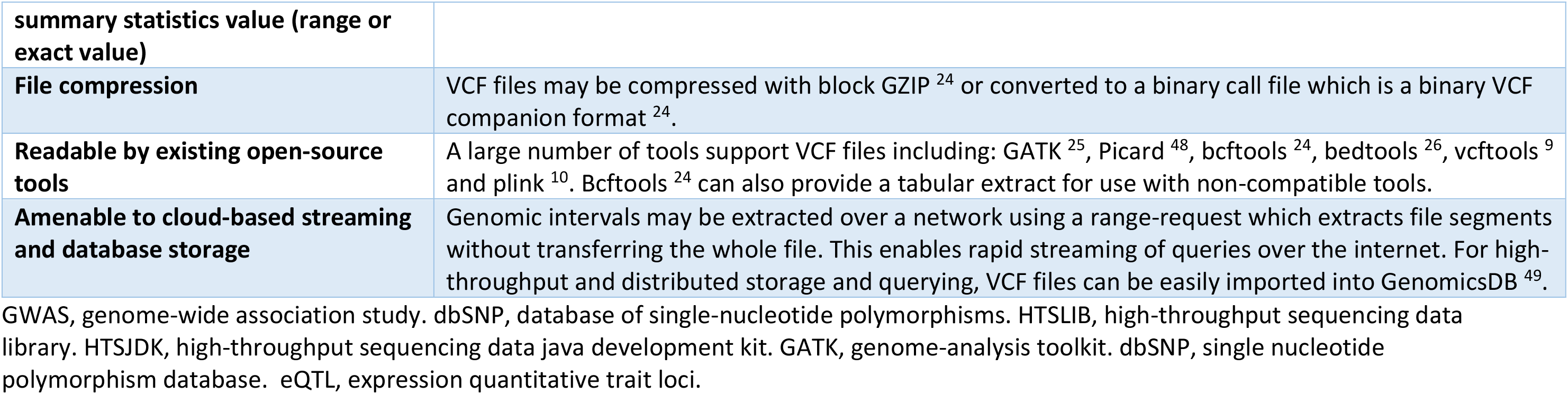
Requirements for a summary statistics storage format and solutions offered by the VCF

The VCF is organised into three components: a flexible file header containing metadata (lines beginning with ‘#’), and a file body containing variant-(one locus per row with one or more alternative alleles/variants) and sample-level information (one sample per column). We adapt this format to include GWAS-specific metadata and utilise the sample column to store variant-trait association data (Figure 1; Supplementary Table 1).

**Figure 1.**
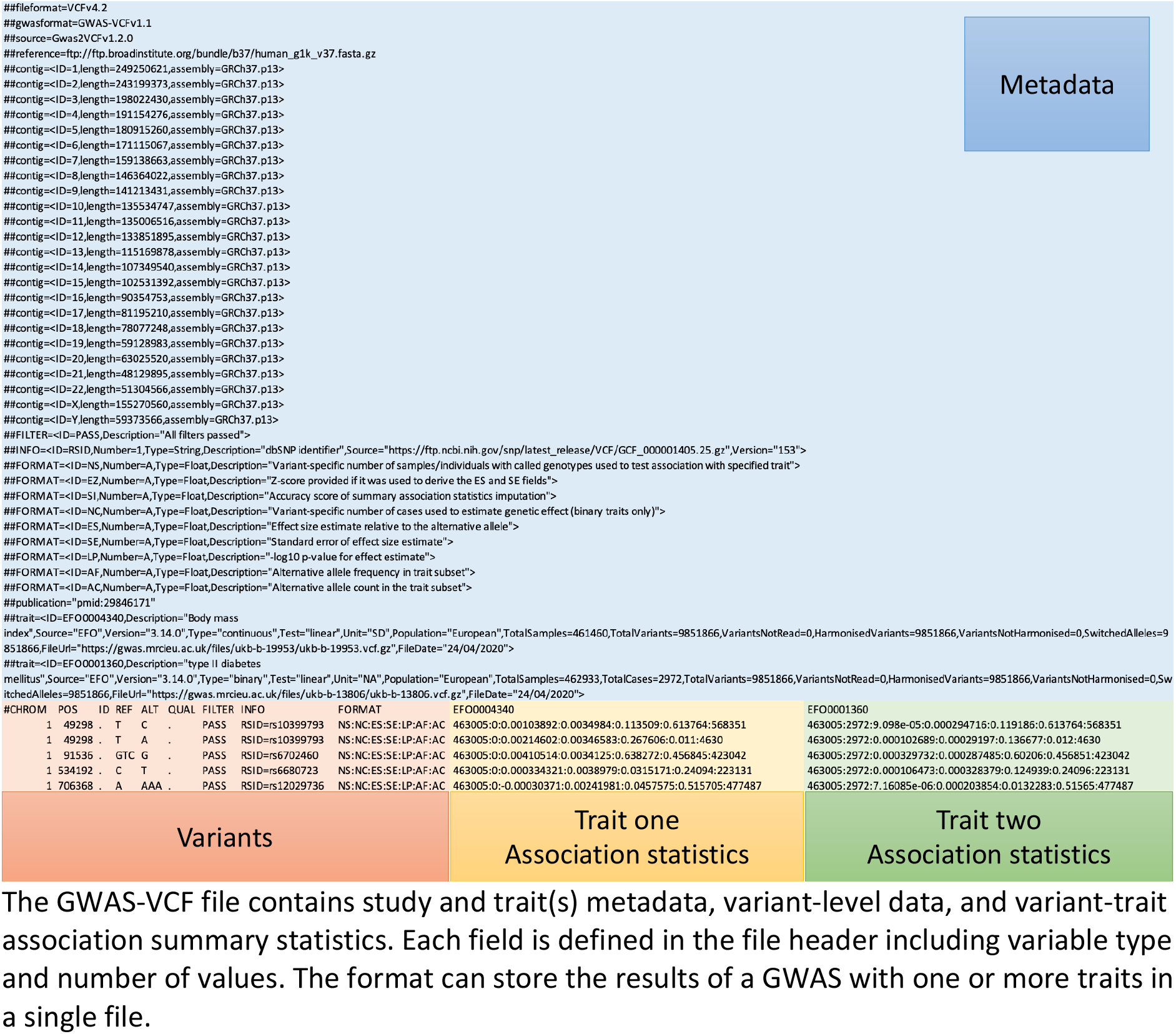
VCF format adapted to store GWAS summary statistics (GWAS-VCF)

**Figure 2.**
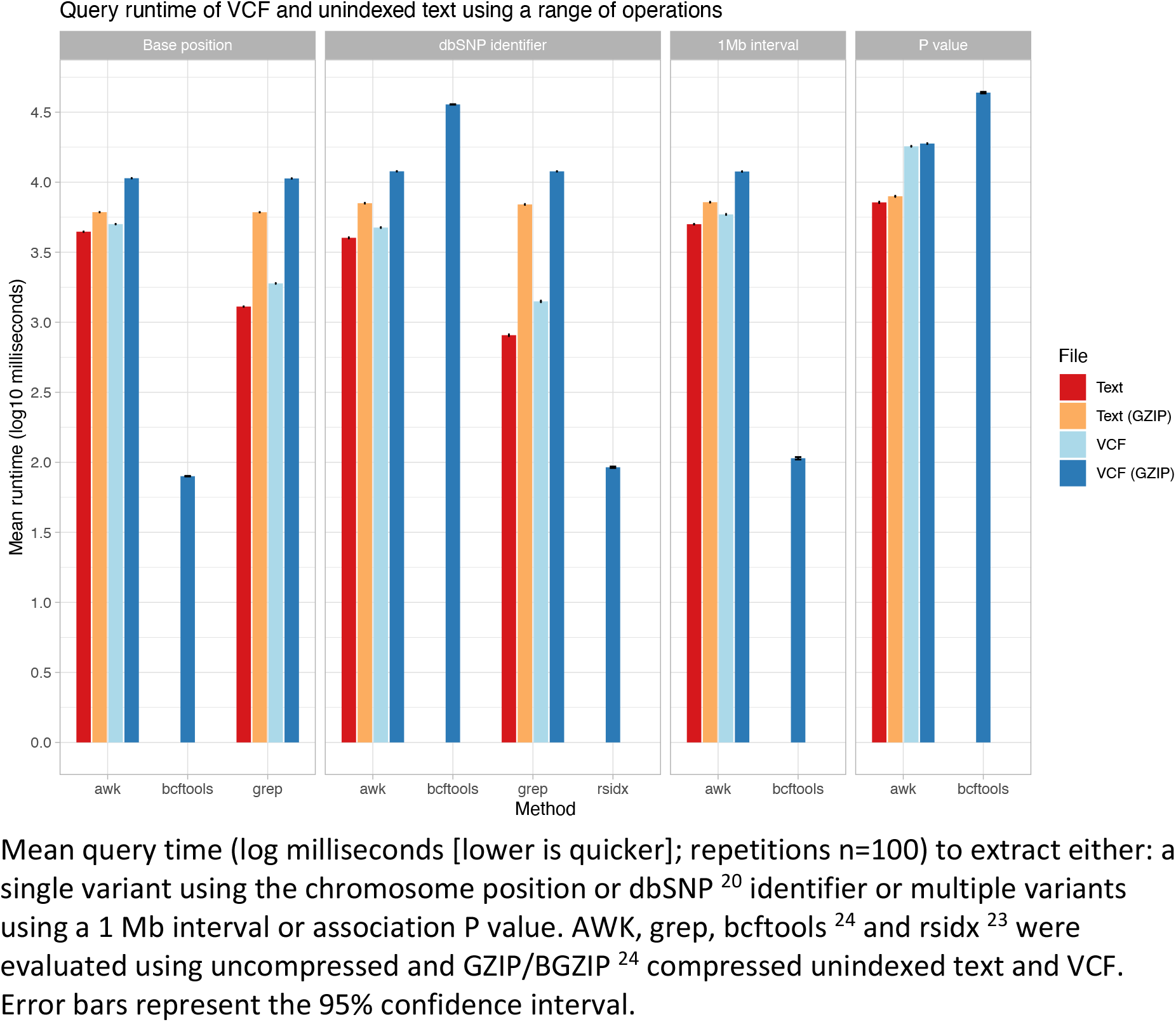
Performance comparison for querying summary statistics in plain text and GWAS-VCF Mean query time (log milliseconds [lower is quicker]; repetitions n=100) to extract either: a single variant using the chromosome position or dbSNP ^20^ identifier or multiple variants using a 1 Mb interval or association P value. AWK, grep, bcftools ^24^ and rsidx ^23^ were evaluated using uncompressed and GZIP/BGZIP ^24^ compressed unindexed text and VCF. Error bars represent the 95% confidence interval.

According to the VCF specification, the file header consists of metadata lines containing 1) the specification version number, 2) information about the reference genome assembly and contigs, and 3) information (ID, number, type, description, source and version) about the fields used to describe variants and samples (or variant-trait associations in the case of GWAS-VCF) in the file body. We take advantage of the VCF file header to store additional information about the GWAS including 1) source and date of summary statistics, 2) study IDs (e.g., PMID/DOI of publication describing the study, or accession number and repository of individual-level data), 3) description of the trait(s) studied (e.g., type, association test used, sample size, ancestry and measurement unit) as well as the source and version of trait IDs (e.g., Experimental Factor Ontology ^17^, Human Phenotyping Ontology ^18^ or Medical Subject Headings ^19^ IDs for clinical and other traits, or Ensembl Gene IDs for eQTL datasets).

Unlike VCF where a row can contain information about multiple alternative alleles observed at the same site/locus (and thus may store more than one variant), the GWAS-VCF specification requires that each variant is stored in a separate row of the file body. Each row contains eight mandatory fields: chromosome name (CHROM), base-pair position (POS), unique variant identifier (ID), reference/non-effect allele (REF), alternative/effect allele (ALT), quality (QUAL), filter (FILTER) and variant information (INFO). The ID, QUAL and FILTER fields can contain a null value represented by a dot. Importantly, the ID value (unless null) should not be present in more than one row. The FILTER field may be used to flag poor quality variants for exclusion in downstream analyses. The INFO column is a flexible data store for additional variant-level key-value pairs (fields) and may be used to store for example: population frequency (AF), genomic annotations and variant functional effects. We also use the INFO field to store the dbSNP ^20^ locus identifier (rsid) for the site at which the variant resides. This is because (despite their common usage as variant identifiers) rsids uniquely identify loci (not variants!) and thus cannot be used in the ID field, as we will discuss further at the end of this manuscript. Following the INFO column is a format field (FORMAT) and one or more sample columns which we use to store variant-trait association data, with values for the fields listed in the FORMAT column for example: effect size (ES), standard error (SE) and −log10 P-value (LP).

This format has a number of advantages over existing solutions. First, the VCF provides consistent and robust approaches to storing genetic variants, annotations and metadata. Furthermore, variable type and number requirements reduce parsing errors and missing data and prevent unexpected program operation. Second, the VCF is well established and supported by existing tools providing a range of functions for querying, annotating, transforming and analysing genetic data. Third, the GWAS-VCF file header stores comprehensive metadata about the GWAS. Fourth, a GWAS-VCF file can store individual or multiple traits (in one or more sample columns) in a single file which is beneficial for the distribution of GWAS datasets where genotypes of each sample/individual have been tested for association with multiple traits (e.g., eQTL datasets).

Simulations of query performance demonstrate compressed GWAS-VCF is substantially quicker than unindexed and uncompressed TSV format for querying by genomic position. On average GWAS-VCF was 16x faster to extract a single variant using chromosome position (mean query duration in GWAS-VCF 0.08 seconds [95% CI 0.08, 0.08]) vs mean query duration in TSV 1.29 seconds [95% CI 1.29, 1.30]) and 9x quicker using the rsid (0.09 seconds [95% CI 0.09, 0.09] vs 0.81 seconds [95% 0.80, 0.82]). Using a 1Mb window of variants GWAS-VCF was 46x quicker (0.11 seconds [95% CI 0.11, 0.11] vs 5.02 seconds [95% CI 4.99, 5.04]). Although querying on association P value was faster using TSV (mean query duration in TSV 7.18 seconds [95% CI 7.09, 7.26] vs mean query duration in GWAS-VCF 18.04 seconds [95% CI 17.92, 18.16]) GWAS-VCF could be improved by using variant flags (i.e. in the INFO field) to highlight records below prespecified thresholds if the exact value is unimportant. For example, all variants below genome-wide significance (P < 5e-8) or a more relaxed threshold (e.g. P < 5e-5).

To automate the conversion of existing summary statistics files to the GWAS-VCF format, we developed open-source Python3 software (Gwas2VCF; Table 2). The application reads in metadata and variant-trait association data using a user-defined schema. During processing, variants are harmonised using a supplied reference genome file to ensure the non-effect allele matches the reference sequence enabling consistent directionality of allelic effects across studies. Insertion-deletion variants are left-aligned and trimmed for consistent representation using the vgraph library ^21^. Finally, the GWAS-VCF is indexed using tabix ^22^ and rsidx ^23^ which enable rapid queries by genomic position and rsid, respectively. We have developed a freely available web application providing a user-friendly interface for this implementation and encourage other centres to deploy their own instance (Table 2).

**Table 2.** Open-source tools for working with GWAS-VCF

| Program | Purpose | Implementation | Source code link |
| --- | --- | --- | --- |
| **gwas2vcf** | Mapping tabular GWAS summary statistics and NHGRI-EBI format to VCF | Python3 (Docker) | https://github.com/mrcieu/gwas2vcf |
| **gwas2vcfweb**<br>http://vcf.mrcieu.ac.uk | Front-end and queue scheduler for gwas2vcf | Python3, Cromwell <sup>50</sup> (Docker) | https://github.com/mrcieu/gwas2vcfweb |
| **R/gwasvcf** | Library for querying and reading GWAS-VCF files | R | https://github.com/mrcieu/gwasvcf |
| **pygwasvcf** | Library for querying and reading GWAS-VCF files | Python3 | https://github.com/mrcieu/pygwasvcf |
| **R/gwasglue** | Library for processing GWAS summary statistics ready for secondary analysis | R | https://github.com/mrcieu/gwasglue |
| **LD Score Regression <sup>5</sup> (patch)** | Estimating genetic correlation and heritability | Python | http://github.com/explodecomputer/ldsc |

GWAS, genome-wide association study. LD, linkage disequilibrium. VCF, variant call format. NHGRI-EBI, National Human Genome Research Institute and European Bioinformatics Institute.

Once stored in a GWAS-VCF file, summary statistics can be read and queried using R or Python programming languages with our open-source libraries (Table 2) or from the command line using for example: bcftools ^24^, GATK ^25^ or bedtools ^26^. Alternatively, GWAS-VCF may be converted to NHGRI-EBI format ^27^ or any other tabular format to support incompatible tools. Further, the gwasglue R package provides convenient programming functions to automate preparation of genetic association data for a range of downstream analyses (Table 2). Currently, methods exist for streamlining variant fine-mapping ^28–32^, colocalization ^33^, MR ^34^ and data visualisation ^35^. New methods are being actively added and users may request new features via the repository issues page.

To encourage adoption, we made openly available over 10,000 complete GWAS summary statistics in GWAS-VCF format as part of the IEU OpenGWAS database. These studies include a broad range of traits, diseases and molecular phenotypes building on the initial collection for the MR Base platform ^34^.

A limitation of current summary statistics formats, including GWAS-VCF, is the lack of a widely adopted and stable representation of sequence variants that can be used as universal unique identifier for said variants. Published summary statistics often use rsids ^20^ to identify variants but this practice is inappropriate because rsids are locus identifiers and do not distinguish between multiple alternative alleles observed at the same site. Moreover, rsids are not stable as they can be merged and retired over time. The reason this is a problem is that in GWAS summary statistics every record represents the effect of a specific allele on one or more traits, and if a record identifier is used that is not unique for each allelic substitution it cannot technically be considered an identifier. An alternative approach is to concatenate chromosome, base-position, reference and alternative allele field values into a single string, but this is non-standardised, and genome build specific. Worst still is the common approach of mixing these types of identifiers within a single file. In version 1.1 of the GWAS-VCF specification we suggest querying variants by chromosome and base-position and filtering the output to retain the target substitution (implemented in our parsers), but we acknowledge that this approach can be cumbersome and difficult to interoperate with other software. The ideal solution would be to populate the ID column of a GWAS-VCF file using universally accepted and unique variant identifiers. We have reviewed several existing variant identifier formats as candidates for the variant identifier field, to be implemented in the next version of the specification (Supplementary Table 2). However, we refrain from making a unilateral choice at this juncture because successful implementation will require consultation from a range of stakeholders. The genetics community uses different approaches already to deal with the problem of sequence variant representation and there is a need to coalesce upon a single format.

Here we present an adaptation of the VCF specification for GWAS summary statistics storage that is amenable to high-throughput analyses and robust data sharing and integration. We implement open-source tools to convert existing summary statistics formats to GWAS-VCF, and libraries for reading or querying this format and integrating with existing analysis tools. Finally, we provide complete GWAS summary statistics for over 10,000 traits in GWAS-VCF. These resources enable convenient and efficient secondary analyses of GWAS summary statistics and support future tool development.

## Code availability

Open-source query performance evaluation source code available from GitHub (https://github.com/MRCIEU/gwas-vcf-performance) or pre-built image available from DockerHub (mrcieu/gwas-vcf-performance)

## Data availability

Version 1.1 of the GWAS - VCF format specification is available from: https://github.com/MRCIEU/gwas-vcf-spec/releases/tag/1.1

Full summary statistics for over 10,000 GWAS in VCF format are available from the IEU OpenGWAS Database (https://gwas.mrcieu.ac.uk)

## Method

### Specification

The specification was developed through experience of collecting and harmonising GWAS summary data across two research centres at scale ^34^ and performing a range of representative high throughput analyses on these data (for example LD score regression ^36^, MR ^37^, genetic colocalisation analysis ^38^ and polygenic risk scores ^39^).

### Query performance simulation

Densely imputed summary statistics (13,791,467 variants) for a large GWAS of body mass index data were obtained from Neale et al ^40^. The data were mapped to VCF using Gwas2VCF v1.1.1 and processed using bcftools v1.10 ^24^ to remove multiallelic variants or records with missing dbSNP ^20^ identifiers. A tabular (unindexed) file was prepared from the VCF to replicate a typical storage medium currently used for distributing summary statistics. Query runtime performance was compared between tabix v1.10.2 ^22^ and standard UNIX commands under the following conditions: single variant selection using dbSNP identifier ^20^ or chromosome position, multi-variant selection by association P value (thresholds: P < 5e-8, 0.2, 0.4, 0.6, 0.8) or 1 Mb genomic interval. Tests were undertaken with 100 repetitions using VCF or unindexed text formats with and without GZIP compression on an Ubuntu v18.04 server with Intel Xeon(R) 2.0 Ghz processor. All comparisons were performed using singled thread operations and therefore differences in runtime performance were due to tool and/or file index usage.

## Acknowledgments

This study was funded by the NIHR Biomedical Research Centre at University Hospitals Bristol National Health Service Foundation Trust and the University of Bristol. The views expressed are those of the author(s) and not necessarily those of the NIHR or the Department of Health and Social Care.

M.L., B.E., T.R.G. work in the Medical Research Council Integrative Epidemiology Unit at the University of Bristol, which is supported by the Medical Research Council and the University of Bristol (MC_UU_00011/4). G.H. is supported by the Wellcome Trust and Royal Society [208806/Z/17/Z].

E.M. and S.J.A. are supported by the JPB foundation and by the National Institute of Health (U01AG052411 and U01AG058635; principal investigator Alison Goate).

## Author contributions

All authors contributed the manuscript and storage format specification. G.H. and E.M. designed the research. M.L. and G.H. wrote software packages and performed query performance simulations. B.E. and G.H. prepared the GWAS data.

---

**Supplementary Table 1.**
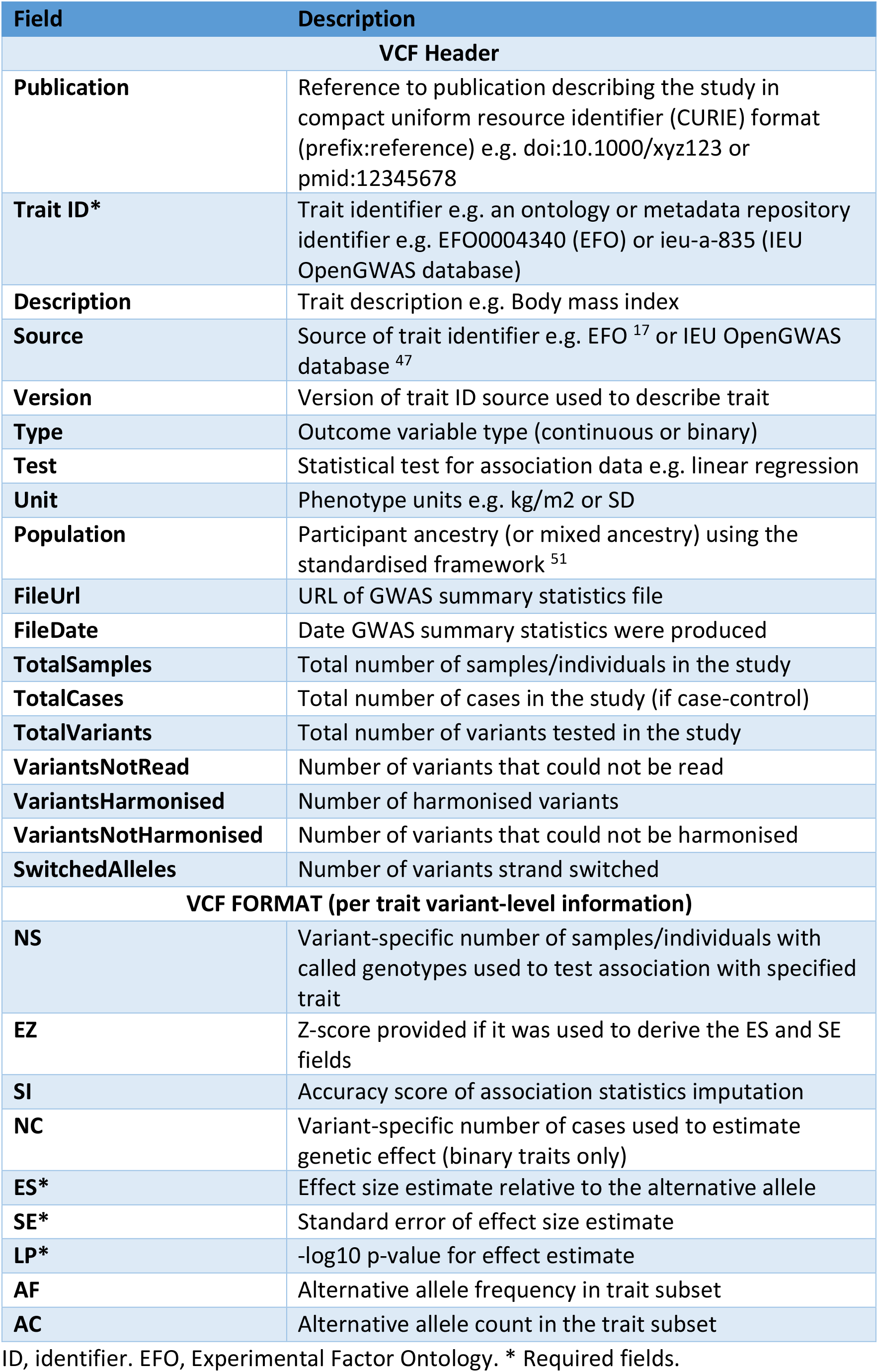
Data fields in the GWAS-VCF

**Supplementary Table 2.**
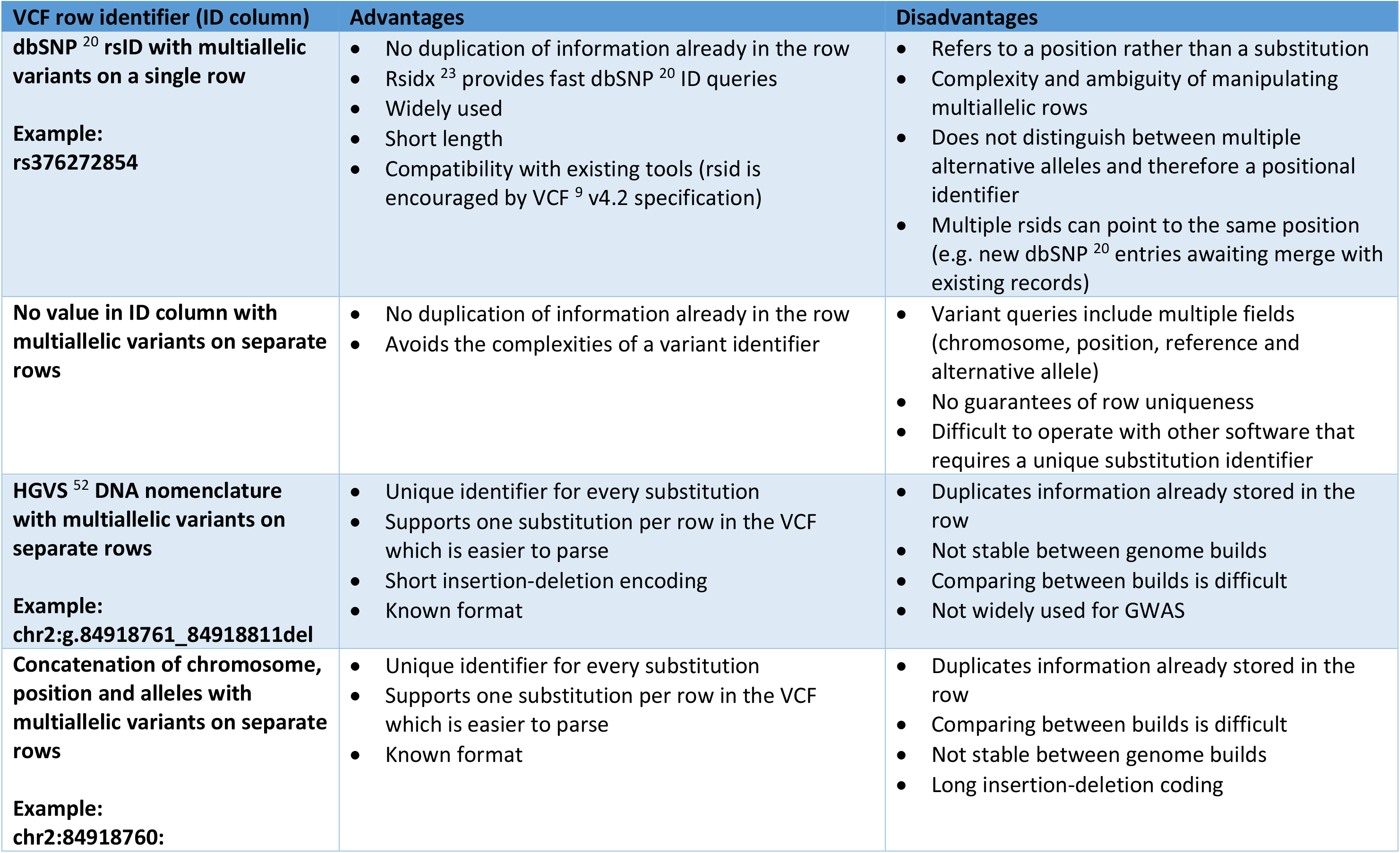
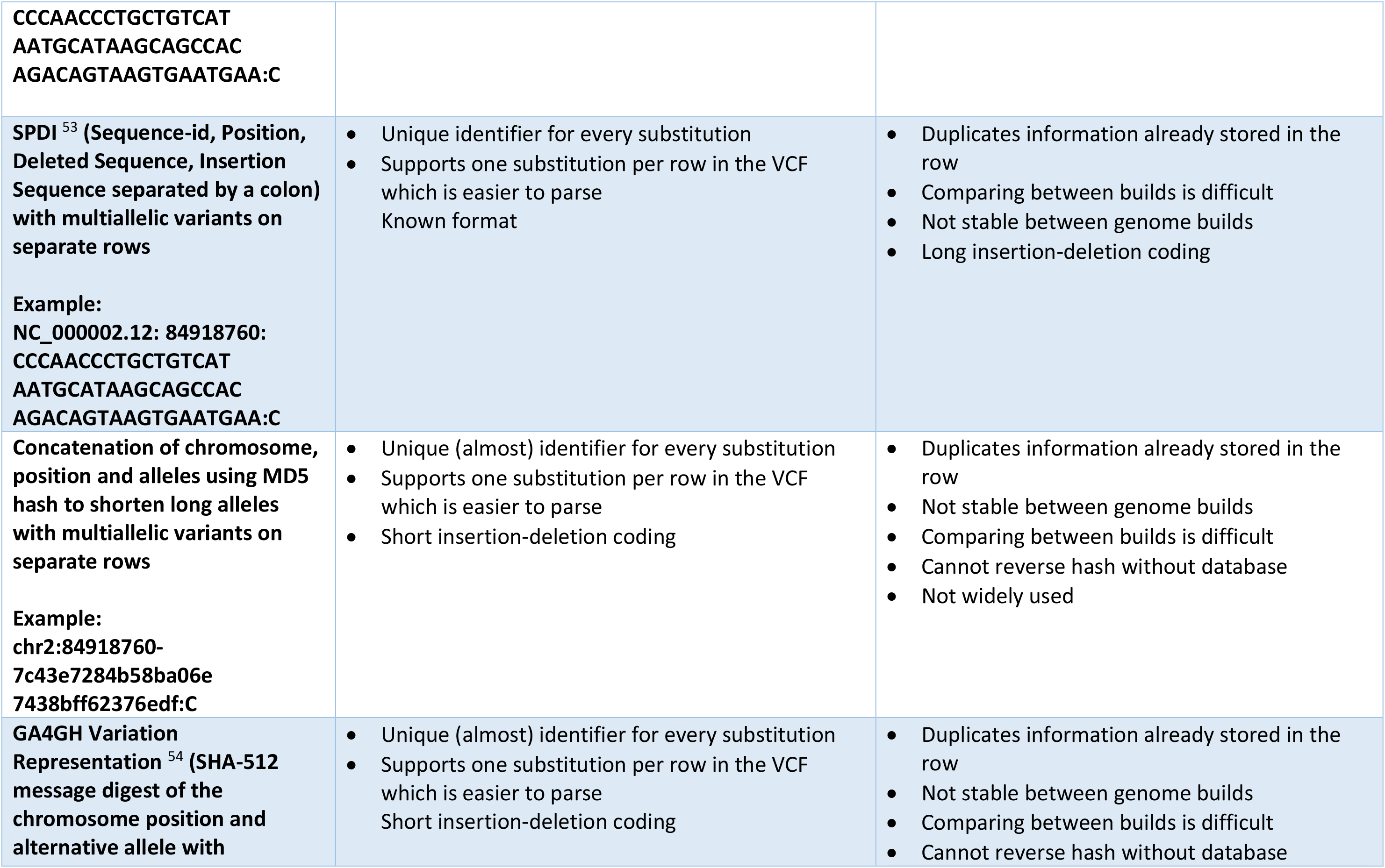
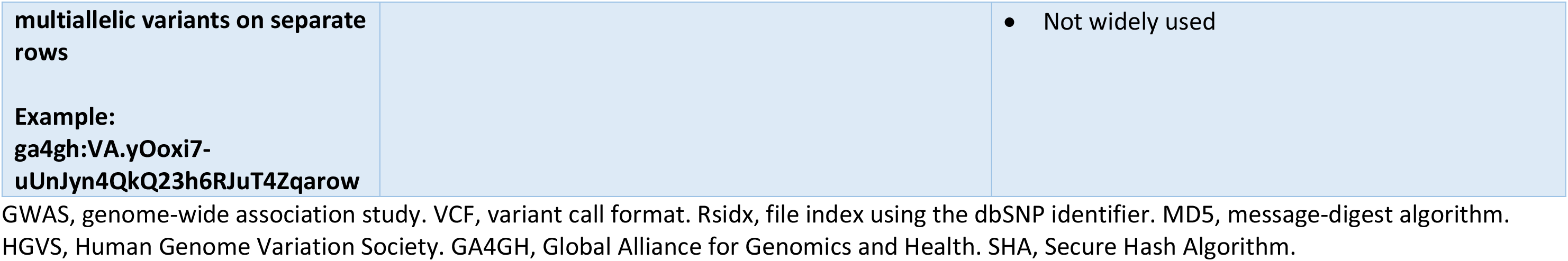
Possible variant identifier schemes for the ID column of GWAS-VCF

